# Predictive coding of visual feedback of actions in human parieto-frontal circuits

**DOI:** 10.64898/2026.09.25.754383

**Authors:** Michelangelo Tani, Sara Ferruzzi, Raffaele Costanzo, Martina Perrone, Gaspare Galati

## Abstract

Executing visually guided movements requires integrating motor commands with visual feedback of the moving hand. Yet, how the motor and visual systems interact during movement remains unquantified. Using real-time motion tracking in immersive virtual reality during fMRI (MOTUM), we manipulated motor execution and hand visual feedback orthogonally in 24 human participants performing reach-to-grasp actions. Execution and hand visual feedback converged in a bilateral parieto-frontal network, including the anterior intraparietal sulcus (aIPS), anterior superior parietal lobule (aSPL), and premotor cortex. In the left aIPS extending into aSPL, combining movement with visual feedback elicited a subadditive response—less activity than predicted by the sum of movement and visual feedback alone—revealing circuitry tuned to the predicted visual consequences of action. Conversely, the posterior cerebellum showed a superadditive response, consistent with internal model updating. Dynamic causal modeling demonstrated a dynamic reversal in information flow along the dorsal parieto-frontal pathway (aSPL–PMd): effective connectivity was excitatory from parietal to frontal cortex when the moving hand was visible, but reversed from frontal to parietal cortex when visual feedback was removed. These findings show that the human parieto-frontal network integrates action and visual feedback predictively, suppressing activity when feedback matches internal predictions and dynamically reconfiguring information flow based on visual availability.

## 1. Introduction

Sensory systems evolved contingent on motor capabilities, with movement being a primary source of evolutionary fitness (Spencer, 1864; Krakauer et al., 2017; Cisek, 2019; Cisek and Hayden, 2022; Cisek and Green, 2024), and sensory feedback serving to refine behavior and increase the precision of action (Cisek, 2007; Pezzulo and Cisek, 2016). Reach-to-grasp actions are the model of choice for studying visuomotor integration, where an object’s visual properties and the sight of one’s own moving hand integrate with proprioceptive signals and are continuously transformed into motor commands (Jeannerod, 1981; Fagg and Arbib, 1998).

Predictive coding provides an operational framework for motor control, with sensorimotor integration rooted in the cross-talk between expectations and prediction errors (Kawato et al., 1987; Wolpert and Kawato, 1998; Wolpert and Flanagan, 2001; Friston, 2010). The inverse internal model computes the difference between current and desired limb position and instantiates the movement. The forward internal model receives a copy of the motor command, predicts its sensory consequences, and compares that prediction with incoming sensory information, eventually signaling the mismatch to the inverse model to adjust the movement. Three consequences follow. (1) When the prediction matches the signal, the response to the two together should be subadditive, that is, below the sum of the responses each evokes alone. This would highlight the circuitry tuned to the expected sensorimotor contingencies we learned from everyday life, such as the seen hand following the motor command. (2) When that prediction is instead being calibrated, the response to the motor and sensory signals together should be superadditive, due to the additional costs of updating the model about novel sensorimotor contingency rules. (3) Finally, since information flows directionally between bottom-up perceptual and top-down motor information, the coupling between the region issuing the prediction, in premotor cortex, and the one testing it, in parietal cortex, should change with the availability of that signal.

While many human neuroimaging studies have contrasted execution with the observation of another person’s hand, a pantomime or a two-dimensional video, or focused on action observation without movement (Culham et al., 2003; Przybylski and Króliczak, 2017; Sulpizio et al., 2020; Bencivenga et al., 2021), testing these hypotheses requires movement and its visual feedback to be manipulated orthogonally within the same participants and session. In virtual reality, the rendered hand can be left at rest while the participant moves, as when reaching in the dark, with the rest of the scene unchanged. Conversely, it can be animated by kinematics recorded earlier while the participant remains still, so that what is seen remains unchanged and only execution is withheld. Moreover, virtual targets can be manipulated so their location and timing cannot be anticipated, making the task engaging and increasing evoked responses, an effect attributed to prediction error (den Ouden et al., 2012).

Here, we leverage MOTUM (Motion Online Tracking Under MRI; Bencivenga and Tani et al., 2026), a novel system that allows real-time kinematics data to be streamed into an immersive virtual environment during fMRI, to let human participants reach and grasp freely while a first-person virtual hand follows (or not) their movements. A factorial design crossing movement and visual feedback yields four conditions differing in execution, in the visibility of the moving hand, and in nothing else, and can answer what contrasting execution with observation cannot. The main effects characterize the large-scale brain networks responding to movement and to hand visual feedback, whether or not the other is present, and their intersection where the two converge. The interaction tests the first two hypotheses: a subadditive response where the motor command already predicts the sight of the hand, and a superadditive response supporting internal model updating. Modulations of the causal effects of the posterior parietal cortex on the premotor cortex, and vice versa (a.k.a. effective connectivity), test the third hypothesis by asking whether their excitatory and inhibitory influences change when the hand becomes visible, as predictive accounts require (Limanowski and Friston, 2020).

## 2. Materials and Methods

### 2.1 Participants

Twenty-four volunteers participated in the study (15 females, mean age 26.6 ± 3.2 years). Participants were right-handed, as assessed by the Edinburgh Handedness Inventory (Oldfield, 1971), had normal or corrected-to-normal vision, and had no neurological or psychiatric disorder. The sample size was based on published recommendations indicating that 20–25 participants provide adequate power to detect medium-to-large effects in novel fMRI paradigms (Desmond and Glover, 2002; Murphy and Garavan, 2004). Participants provided written informed consent, and the regional research ethics committee approved the study (Prot. CE/PROG.932).

### 2.2 Experimental Design

The experiment manipulated motor execution and visual feedback from the moving hand orthogonally within the same participants and the same session. We used a 2×2 factorial design with movement (executed or not) and hand visual feedback (present or absent) as factors, yielding four conditions: Move Visible (MV), Move Invisible (MI), Stay Visible (SV), and Stay Invisible (SI).

Each condition isolates a different aspect of visually guided reach-to-grasp. MV is the complete action, executed with online vision of a virtual hand in first-person perspective, animated in real time by the participant’s movements. MI removes visual feedback while preserving execution, so the reach is guided by proprioception, as it is when reaching in the dark. SV preserves the sight of hand motion without execution, the virtual hand being animated by the kinematics of movements the same participant executed seconds earlier on MV trials, as a 3D video replay of the movement. SI removes both the task and the stimulus and retains only the target cue, with the virtual arm again at rest, thereby serving as the control condition against which the other three are expressed.

Replay makes the visual feedback factor interpretable. Contrasting execution with observing another person’s hand, a pantomime, or a two-dimensional video changes the hand’s ownership, kinematics, and identity, as well as the low-level features of the visual feedback. Because the kinematics rendered on the virtual hand are the participant’s own and are rendered as they were produced, MV and SV differ in execution and in nothing else that the participant sees, and SV and SI differ in the sight of one’s own moving hand and in nothing else.

### 2.3 Stimuli and tasks

Throughout the experiment, participants lay supine on the MR scanner bed and viewed a virtual environment simulating a scanner room (Figure 1) through a binocular headset. Participants saw a humanoid avatar’s arm in a first-person perspective. A target object floated above a wooden table and consisted of a custom symmetric 3D sinusoidal shape with four alternating wide and narrow sections. In each trial, one of the four central sections of the virtual object was randomly illuminated for 2.5 s. The wide sections afforded a whole-hand power grip and the narrow ones a precision grip between the thumb and index finger, so neither the grip nor its location could be anticipated, which kept the task engaging.

**Figure 1.**
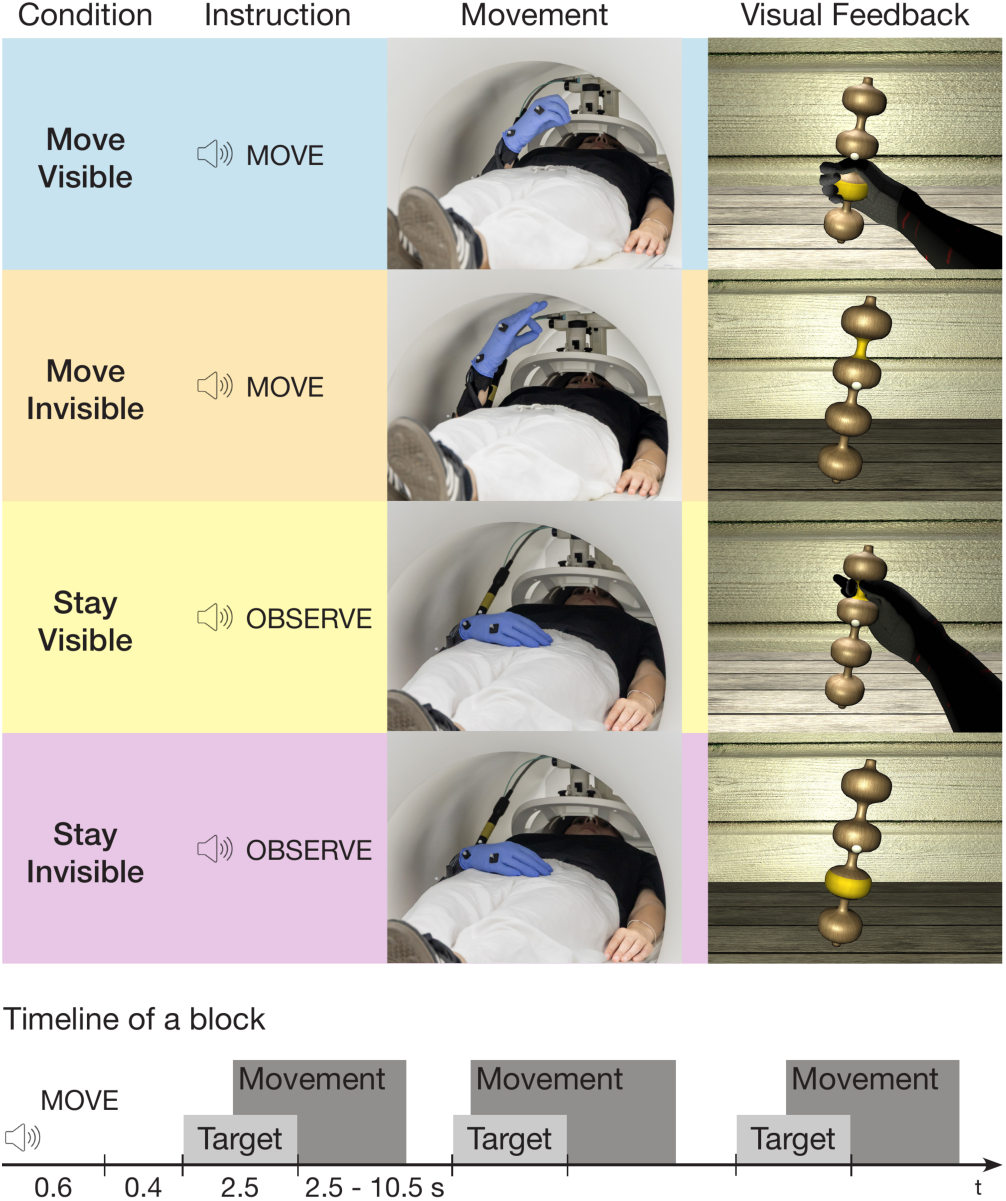
Top: the 2×2 factorial design with movement execution (Move/Stay) and hand visual feedback (Visible/Invisible), yielding four conditions: Move Visible, Move Invisible, Stay Visible, and Stay Invisible. Each condition shows the instruction type, participant posture (moving or at rest), and the corresponding visual feedback provided through the goggles. Bottom: timeline of a single experimental block. Each block began with an auditory instruction ("Move" or "Observe"), followed by three trials. Each trial started with a yellow light cue lasting 2.5 s, marking one of the four central sections of the virtual object: either a narrow section requiring a precision grip or a wide section requiring a power grip. The intertrial interval was jittered (range 2.5–10.5 s, mean 4.5 s).

Conditions were delivered in short blocks, preceded by an auditory instruction (“Move” or “Observe”, 1 s) to indicate to the participant whether to move or to remain still on the trials that followed. Participants were asked to maintain fixation on a white sphere at the virtual object’s center for the whole experiment, and to keep their right palm positioned comfortably on their abdomen before and after each trial.

When instructed to “Move”, participants reached and grasped the illuminated section and then returned to the starting position. In Move Visible (MV) blocks, the virtual hand was animated in real time by the participant’s movements, providing online visual feedback. In contrast, in Move Invisible (MI) blocks, the virtual hand did not follow the movement and remained on the virtual table in a resting posture; therefore, participants removed the sight of the hand moving, not the hand’s presence in the scene.

When instructed to “Observe”, participants remained still. In Stay Visible (SV) blocks, the virtual hand was animated using the kinematics of randomly selected previous MV trials, along with the cue that elicited them. Hence, the movement always matched the cued section, and participants saw their own hand kinematics without executing a movement. In Stay Invisible (SI) blocks, the virtual hand likewise remained at rest, and only the cue appeared, so the visual scene was identical in MI and SI and differed from MV and SV only in hand motion (Figure 1).

Trials lasted 2.5 s and were separated by jittered intervals drawn from an exponential distribution with a mean of 4.5 s (range 2.5–10.5 s), allowing us to model and estimate responses for each trial separately. The experiment included four fMR scans, each consisting of 20 blocks arranged into five sets, each containing all four block types. Block order within a set was assigned pseudorandomly to maximize heterogeneity, considering only permutations in which SV blocks did not appear before MV blocks, so that recorded kinematics were available for rendering. No block order was used more than twice per session, out of the 12 permutations that satisfied this constraint. Grip type (precision or power) and grip location (top, mid-top, mid-bottom or bottom central sections of the virtual object) were randomly distributed and counterbalanced within runs across the executed conditions. Each scan lasted approximately 7 min 30 s, and the whole session lasted approximately 80 min including participant preparation and structural imaging.

### 2.4 Apparatus and procedure

We used MOTUM (Bencivenga and Tani et al., 2026; Figure 2) installed at the Neuroimaging Laboratory, LIFE Institute for Research and Health Care Santa Lucia, Rome, Italy, with a Magnetom Prisma 3 T scanner (Siemens, Erlangen, Germany) and a standard 64-channel head coil. Motion capture was achieved through seven electromagnetically shielded cameras (three Oqus 500 MRI and four Arqus MRI; Qualisys, Göteborg, Sweden) positioned on the frontal and lateral walls of the scanner room to track hand and forearm position, complemented by an MR-compatible kinematic glove (Data Glove Ultra, Fifth Dimension Technologies, Pretoria, South Africa) equipped with 14 sensors for capturing finger flexion/extension. The virtual environment was created in Unity 6 (Unity Technologies, San Francisco, USA), which received motion data from Qualisys Track Manager via TCP/IP and glove data via serial connection, managed stimulus presentation and timing synchronization, and rendered the scene to high-definition MR-compatible goggles (VisualSystem HD, Nordic NeuroLab, Bergen, Norway; 1920×1080 resolution at 60 Hz). We delivered auditory instructions through MR-compatible headphones with active noise control (OptoActive II, Optoacoustics Ltd, Mazor, Israel), which attenuated scanner noise throughout the session. We maintained scanner-stimulus synchronization via an optical trigger line (FIU 932, Current Designs, Philadelphia, USA). For further details on the MOTUM system, see Bencivenga and Tani et al. (2026).

**Figure 2.**
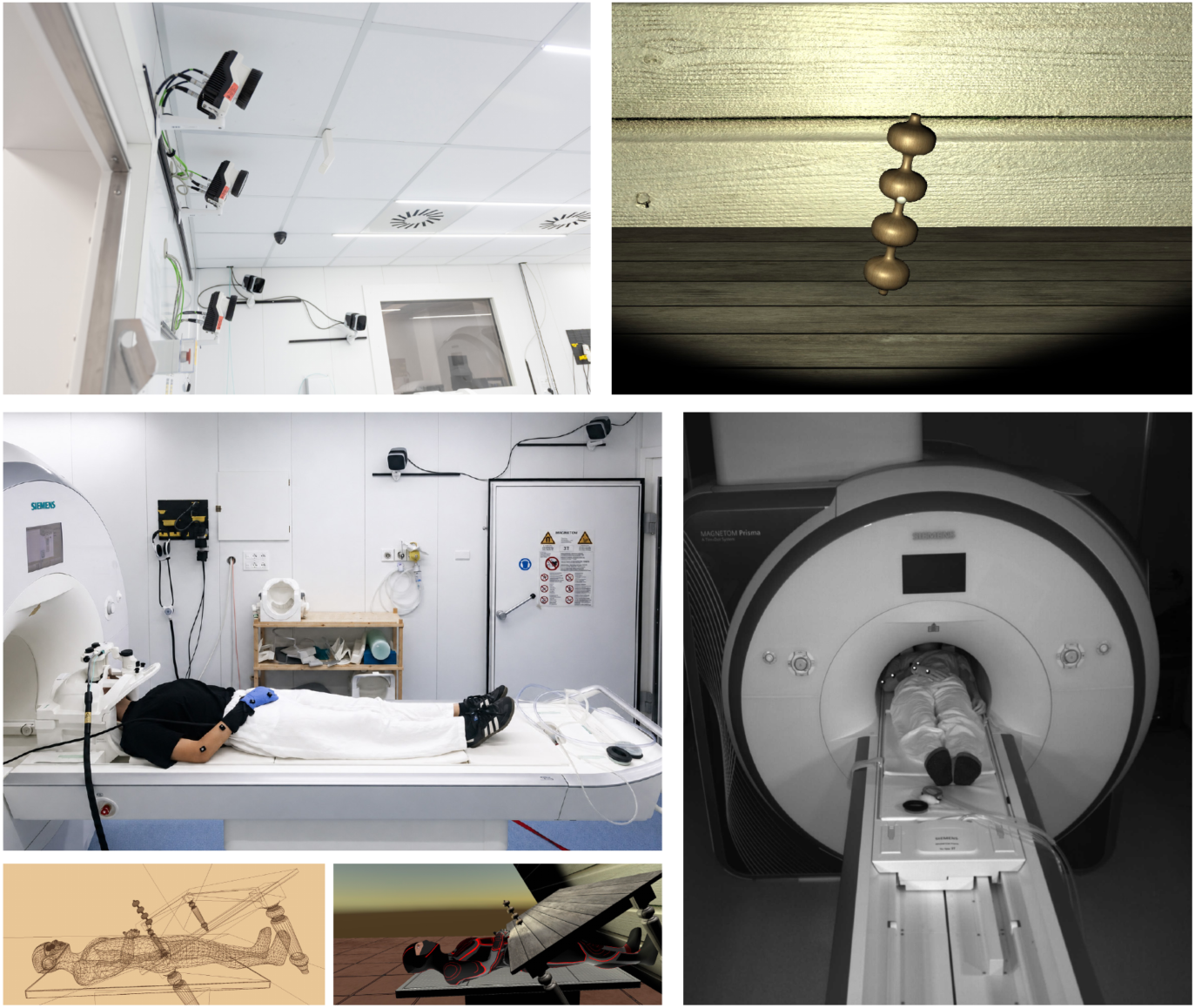
MOTUM, experimental apparatus. Top left: the array of seven electromagnetically shielded motion-tracking cameras (Oqus 500 MRI and Arqus MRI, Qualisys) mounted on the frontal and lateral walls of the scanner room. Top right: the virtual target object, a symmetric sinusoidal shape whose four alternating wide and narrow sections afford a power or a precision grip, with one section illuminated to cue the reach. Mid-left: participant positioned on the scanner table, outside the bore, wearing the MR-compatible virtual reality goggles and kinematic glove, with motion-tracking markers attached to the arm. Bottom-right: view from one of the frontal-wall tracking cameras, showing the participant at the scanner isocenter with visible tracking markers. Bottom-left: wireframe rendering of the virtual environment, showing the polygonal structure of the scene, including the avatar, the table, and the tilted graspable 3D object. Bottom-middle: fully rendered view of the same virtual environment, from the avatar’s first-person point of view.

Before the experimental session, participants completed the informed consent form, were screened by qualified medical personnel, and were instructed about the task and the safety rules. Each session began with camera-system calibration to a room-based right–anterior– superior (RAS) coordinate system (Bencivenga and Tani et al., 2026). After entering the scanner, we fitted the participant’s right hand with the kinematic glove and attached five markers to the right arm and hand. We secured participants’ heads in the head coil with cushions for comfort and stability, along with MR-compatible headphones and goggles. We adjusted the pupillary distance and the distance from the eyes to the goggles for participant comfort and visual clarity. A customized graphical user interface allowed us to adapt the virtual environment to each participant’s range of motion (i.e., the position of the virtual object). Participants then practiced the movements for several minutes to familiarize themselves with the virtual avatar before scanning began. At rest, the participant’s right hand was positioned on their abdomen, whereas in the virtual scene, it appeared to rest on a table in front of them, viewed from a first-person perspective.

During the MR session, we acquired the following scans:

1. One T1-weighted structural image was acquired with a magnetization-prepared rapid gradient-echo (MPRAGE) sequence (176 slices, isotropic resolution 1 mm, field of view 256 × 256 mm², TR 2500 ms, TE 1.67 ms, inversion time 1080 ms, flip angle 8°), with prospective motion correction and selective reacquisition of data corrupted by motion based on interleaved 3D EPI navigators (Hess et al., 2011; Tisdall et al., 2012).
2. One T2-weighted structural image was acquired with a 3D SPACE sequence with variable flip angle refocusing pulses at a nominal flip angle of 120° (176 slices, isotropic resolution 1 mm, field of view 256 × 240 mm², TR 3200 ms, TE 564 ms), and prospective motion correction as above.
3. Four functional T2*-weighted scans were acquired with a gradient-echo echo-planar imaging (EPI) sequence (multiband factor 6, isotropic voxel size 2.4 mm, 60 slices, field of view 211 × 211 mm², TR 800 ms, TE 30 ms, flip angle 52°, no in-plane acceleration) (Feinberg et al., 2010; Moeller et al., 2010; Xu et al., 2013). During scan acquisition, participants completed the experimental tasks using four different pseudorandomized trial lists. Scan duration varied by trial list (mean 561.8 volumes per run, SD 17.8).
4. We acquired two spin-echo EPI volumes for field mapping with phase encoding in opposite directions, no multiband acceleration, and the same geometrical and sampling properties as the functional runs (TR 7060 ms, TE 80 ms).

After the experimental session, participants completed a six-item questionnaire to assess embodiment on a 10-point rating scale (Botvinick and Cohen, 1998; Tieri et al., 2015). Two measures were obtained. Ownership, the degree to which participants felt that the virtual arm belonged to them, was the average of responses to Items 1 and 2, with Item 3 as a control. Agency, the feeling of control over the virtual hand’s actions, was the average of responses to Items 4 and 5, with Item 6 as a control. Participants reported a sense of ownership of 5.04 ± 2.2 and a sense of agency of 6.64 ± 2.2 on the 10-point scale, whereas the two control items, which describe experiences that should not occur, averaged 1.60 ± 1.1.

### 2.5 Data processing

#### 2.5.1 Imaging data processing

Image preprocessing was performed using fMRIPrep 25.0.0 (Esteban et al., 2018, 2019), which is based on Nipype 1.9.2 (Gorgolewski et al., 2011, 2018). The following text was automatically generated by fMRIPrep with the express intention to be copied and pasted into manuscripts unchanged, and is released under the CC0 license.

The T1w image was corrected for intensity non-uniformity (INU) with N4BiasFieldCorrection (Tustison et al., 2010), distributed with ANTs 2.5.4 (Avants et al., 2008), and used as T1w-reference throughout the workflow. The T1w-reference was then skull-stripped with a Nipype implementation of the antsBrainExtraction.sh workflow (from ANTs), using OASIS30ANTs as target template. Brain tissue segmentation of cerebrospinal fluid (CSF), white matter (WM), and gray matter (GM) was performed on the brain-extracted T1w using fast (FSL; Zhang et al., 2001). Brain surfaces were reconstructed using recon-all (FreeSurfer 7.3.2; Dale et al., 1999), and the brain mask estimated previously was refined with a custom variation of the method to reconcile ANTs-derived and FreeSurfer-derived segmentations of the cortical gray matter of Mindboggle (Klein et al., 2017). The T2-weighted image was used to improve pial surface refinement. Volume-based spatial normalization to two standard spaces (MNI152NLin6Asym, MNI152NLin2009cAsym) was performed through nonlinear registration with antsRegistration (ANTs 2.5.4), using brain-extracted versions of both T1w reference and the T1w template. The following templates were selected for spatial normalization and accessed with TemplateFlow (24.2.2; Ciric et al., 2022): FSL’s MNI ICBM 152 non-linear 6th Generation Asymmetric Average Brain Stereotaxic Registration Model (Evans et al., 2012; TemplateFlow ID: MNI152NLin6Asym), ICBM 152 Nonlinear Asymmetrical template version 2009c (Fonov et al., 2009; TemplateFlow ID: MNI152NLin2009cAsym). Grayordinate “dscalar” files containing 91k samples were resampled onto the fsLR (fs_LR 32k) surface mesh using the Connectome Workbench (Glasser et al., 2013).

For each of the blood oxygenation level-dependent (BOLD) runs, the following preprocessing was performed. First, a reference volume was generated using a custom methodology of fMRIPrep for head motion correction. Head-motion regressors with respect to the BOLD reference (transformation matrices and six corresponding rotation and translation parameters) are estimated before any spatiotemporal filtering using mcflirt (FSL; Jenkinson et al., 2002). The estimated fieldmap was then aligned with rigid registration to the target EPI reference run. The field coefficients were mapped onto the reference EPI using the transform. The BOLD reference was then co-registered to the T1w reference using bbregister (FreeSurfer), which implements boundary-based registration (Greve and Fischl, 2009). Co-registration was configured with six degrees of freedom. The aligned T2w image was used for initial co-registration. Several confounding time series were calculated based on the preprocessed BOLD: framewise displacement (FD), the derivative of root mean square variance over voxels (DVARS), and three region-wise global signals. FD was computed using two formulations following Power (absolute sum of relative motions; Power et al., 2014) and Jenkinson (relative root mean square displacement between affines; Jenkinson et al., 2002). FD and DVARS are calculated for each functional run, both using their implementations in Nipype (following the definitions by Power et al., 2014). The three global signals are extracted within the CSF, the WM, and the whole-brain masks. Additionally, physiological regressors were extracted to allow for component-based noise correction (CompCor; Behzadi et al., 2007). Principal components are estimated after high-pass filtering the preprocessed BOLD time-series (using a discrete cosine filter with 128s cut-off) for the two CompCor variants: temporal (tCompCor) and anatomical (aCompCor). tCompCor components are then calculated from the top 2% variable voxels within the brain mask. For aCompCor, three probabilistic masks (CSF, WM and combined CSF+WM) are generated in anatomical space. The implementation differs from that of Behzadi et al. in that instead of eroding the masks by 2 pixels on BOLD space, a mask of pixels that likely contain a volume fraction of GM is subtracted from the aCompCor masks. This mask is obtained by dilating a GM mask extracted from the FreeSurfer’s aseg segmentation, and it ensures components are not extracted from voxels containing a minimal fraction of GM.

Finally, these masks are resampled into BOLD space and binarized by thresholding at 0.99 (as in the original implementation). Components are also calculated separately within the WM and CSF masks. For each CompCor decomposition, the k components with the largest singular values are retained, such that the retained components’ time series are sufficient to explain 50 percent of the variance across the nuisance mask (CSF, WM, combined, or temporal). The remaining components are dropped from consideration. The head-motion estimates calculated in the correction step were also placed within the corresponding confounds file. The confound time series derived from head motion estimates and global signals were expanded with the inclusion of temporal derivatives and quadratic terms for each (Satterthwaite et al., 2013). Frames that exceeded a threshold of 0.5 mm FD or 1.5 standardized DVARS were annotated as motion outliers. Additional nuisance timeseries are calculated by means of principal components analysis of the signal found within a thin band (crown) of voxels around the edge of the brain, as proposed by Patriat et al. (2017). The BOLD time series were resampled onto the left/right-symmetric template “fsLR” using the Connectome Workbench (Glasser et al., 2013). A “goodvoxels” mask was applied during volume-to-surface sampling in fsLR space, excluding voxels whose time series have a locally high coefficient of variation. Grayordinates files (Glasser et al., 2013) containing 91k samples were also generated with surface data transformed directly to fsLR space and subcortical data transformed to 2 mm resolution MNI152NLin6Asym space. All resamplings can be performed with a single interpolation step by composing all the pertinent transformations (i.e., head-motion transform matrices, susceptibility distortion correction when available, and co-registrations to anatomical and output spaces). Gridded (volumetric) resamplings were performed using nitransforms, configured with cubic B-spline interpolation. Finally, a spatial smoothing was performed on the cortical surface using geodesic smoothing with a 6 mm full-width at half-maximum (FWHM) kernel (Hagler et al., 2006).

#### 2.5.2 Kinematic data processing

We reconstructed kinematic data from arm and hand movements in real time from the virtual environment and saved them at 60 fps; we analyzed the position and rotation of the wrist, thumb tip, and index finger tip. The kinematics served a single purpose here: to segment each trial into its reaching, grip, and back-movement phases so the duration of the reaching phase could be entered as a parametric modulator in the imaging model. A custom k-means clustering algorithm identified the transitions between phases, and we plotted and visually inspected the kinematics of every trial to detect errors in the participant’s behavior or in the segmentation.

Trials were excluded when movement was absent, incorrect, or repeated in move trials; when the arm moved during a stay condition; when marker tracking was lost; when a software error in an early version of the stimulus presentation package delivered incorrect visual feedback; or when phase segmentation failed. Of the 240 trials planned for each session, we excluded an average of 15 trials on these grounds. We modeled excluded trials as error regressors in the imaging analyses.

### 2.6 Statistical analysis

#### 2.6.1 Activation analyses

We analyzed functional images for each participant in grayordinate space, comprising the fsLR 32k surface vertices (FreeSurfer 7.3.2; Dale et al., 1999; Glasser et al., 2013) and subcortical voxels, using a general linear model implemented in SPM25 (Wellcome Center for Human Neuroimaging, London, UK). We smoothed grayordinate time series with a 6 mm full-width-at-half-maximum Gaussian kernel applied along the cortical surface over geodesic distances. Within the subcortical volume, we high-pass filtered data at 128 s and modeled serial correlations with a first-order autoregressive model. Each trial was modeled as a boxcar function of 2.5 s (cue duration) convolved with the canonical hemodynamic response function, with no time or dispersion derivatives. We included a separate regressor for each trial type: MV, MI, SV, and SI. For each trial type in which the hand moved or was seen to move (MV, MI, and SV), we included an additional regressor modeling reaching-phase duration as a linear parametric modulator. We included error trials with and without movement as separate regressors of no interest.

To control for motion-related artifacts, we included framewise displacement (head motion; Power et al., 2012) and four CompCor components as nuisance regressors, along with a constant term. Because participants moved their arms inside the bore by design, movement itself is the experimental factor, and regressing out movement-related variance beyond head motion would remove signals of interest along with artifacts. We quantified the consequences of arm and hand movement for data quality in this setup separately (Bencivenga and Tani et al., 2026). We therefore used framewise displacement to account for residual head motion. Individual statistical parametric maps were obtained for contrasts testing the main effects of movement ([MV + MI] > [SV + SI]) and of hand visual feedback ([MV + SV] > [MI + SI]), as well as for interaction analyses testing superadditive effects ([MV – MI] > [SV − SI]) and subadditive effects ([MV − MI] < [SV − SI]). We then entered the resulting contrast images into a group-level random-effects analysis, a one-sample t-test across the 24 participants, computed separately for each contrast.

We thresholded statistical maps at *p* < 0.05, corrected for multiple comparisons at the cluster level using topological false discovery rate (FDR) correction based on random field theory (Chumbley et al., 2010), after applying a cluster-forming threshold of *p* < 0.001 uncorrected. We masked interaction maps by activation in either MI or SV relative to SI.

We identified regions responding to both factors as the intersection of grayordinates exceeding the threshold in both main effect maps. This overlap localizes convergence and is not itself a statistical test; we draw no inference from it beyond what each main effect supports separately.

#### 2.6.2 Effective connectivity analyses

To characterize how movement execution and hand visual feedback shape effective connectivity within the reach-to-grasp network, we modeled a network connecting the main nodes of the parieto-frontal circuits that transform visual and somatosensory information into motor commands (see the Results section for details). Because participants moved only their right hand, we restricted the model to intrahemispheric connectivity in the contralateral left hemisphere.

We estimated effective connectivity with dynamic causal modeling (DCM; Friston et al., 2003), which models the hidden neuronal activity generating the BOLD signal through an extended balloon model (Buxton et al., 1998; Friston et al., 2000). A bilinear approximation decomposes these dynamics into three sets of parameters, namely intrinsic (endogenous) connectivity between regions, which holds throughout the session and is the only coupling present when no condition is active (matrix A), condition-specific changes in that connectivity (matrix B), and direct driving inputs onto each region for a given condition (matrix C). Each condition entered the model as the same 2.5 s boxcar used in the first-level model, locked to the cue, so the driving inputs and modulations act over the interval spanning the cue, the reach, the grip, and, on most trials, the onset of the return.

We did not insert SI blocks into the connectivity model. Every trial began with a target cue, and SI trials contained nothing else, so leaving that condition unmodeled treats the cue-evoked response as part of the baseline, together with the block instructions, and so implicitly models them into the intrinsic connectivity component (matrix A), against which the driving inputs and modulations estimated for the other three conditions express what the movement of the hand and the visual feedback of that movement add.

Connectivity nodes included the primary motor cortex (M1) and a set of nodes defined from the intersection of the activation maps of the two main effects, that is, from the grayordinates exceeding threshold both for movement and for hand visual feedback, so that each node lay in tissue responsive to both factors. Matrix A was specified a priori from known cortico-cortical pathways within and across the selected nodes. Driving inputs (matrix C) were restricted to a single node per condition. A condition that both drives several nodes and modulates the connections between them makes the driving and the modulatory parameters partly redundant, since the same variance can be accommodated in either, and keeping the inputs sparse leaves the condition-specific effects to matrix B, where the hypotheses of this study are expressed (Zeidman et al., 2019a). Modulations (matrix B) were specified on a subset of the intrinsic connections. Details of the included nodes, connections, and modulations are described in the Results section, since they critically depended on the activation analysis results.

We derived connectivity nodes from activation maps using watershed segmentation (Mangan and Whitaker, 1999) to select precise local maxima and their neighborhoods within activation clusters spanning multiple regions. Where applicable, nodes were defined from each participant’s own first-level statistical maps. If a participant showed no significant activation at a given node, we substituted the corresponding group-level region, preserving network coverage while retaining participant specificity.

For each node, we extracted a representative time series from unsmoothed, preprocessed data as the first eigenvariate, adjusted for confounds modeled in the general linear model. We submitted these time series to first-level DCM estimation. We inverted one model per participant using the variational Laplace scheme (Friston et al., 2007), applying variational free energy as a proxy for model evidence. We flagged any model explaining less than 10% of the BOLD variance as poorly inverted (Zeidman et al., 2019a). Rather than excluding these participants outright, we carried them forward into the group-level analysis and downweighted their contribution according to posterior certainty.

At the group level, we used Parametric Empirical Bayes (PEB; Friston et al., 2015, 2016) to combine the individual DCMs into a second-level model accounting for between-subject variability (Zeidman et al., 2019b). The design matrix, X = [1…1]ᵀ, modeled only the group mean. We analyzed matrices A, B, and C in three separate PEBs to limit evidence dilution across a large parameter space (Zeidman et al., 2019b). We then applied Bayesian model reduction with greedy search, as implemented in SPM25, to prune model parameters iteratively in batches rather than enumerating all 2ᵏ nested models (Friston and Penny, 2011; Pinotsis et al., 2016), and Bayesian model averaging to average parameters over the retained models, weighted by their evidence (Friston and Penny, 2011). We report parameters when the posterior likelihood that they differ from zero exceeds 90%, the threshold conventionally adopted in group DCM studies as evidence that a parameter is present (Zeidman et al., 2019b). Posterior probability indexes confidence that a parameter is non-zero, not the magnitude of the effect, which is given by the parameter estimate itself, in Hz for connections between regions and as a log-scaling factor for self-connections.

### 2.7 Data and Code Accessibility

We carried out preprocessing with fMRIPrep 25.0.0, whose workflow is described above. We ran all subsequent analyses in MATLAB 2025b with SPM25. The complete dataset, including raw and preprocessed images, kinematic data, and statistical models, will be available on request to the corresponding author, subject to a formal data-sharing agreement and approval by the requesting researcher’s local ethics committee.

## 3. Results

### 3.1 A parieto-frontal network shared by movement and hand visual feedback

Executing reach-to-grasp movements, irrespective of whether the hand was visible (main effect of movement: [MV + MI] > [SV + SI]), engaged a network centered on the sensorimotor representation of the hand and arm contralateral to the moving hand and extending anteriorly into premotor and medial motor cortex and posteriorly into the superior parietal lobule and the intraparietal sulcus (Figure 3, yellow). Further clusters were located bilaterally in the middle frontal gyrus, basal ganglia, thalamus, brainstem, and cerebellum. Seeing one’s own hand move, irrespective of whether the participant moved (main effect of visual feedback: [MV + SV] > [MI + SI]), engaged a network running from early visual and motion-sensitive occipito-temporal cortex, including the human middle temporal complex (hMT+) and the fusiform body area, into the intraparietal sulcus and the superior parietal lobule (Figure 3, blue). Separate clusters were located in the dorsal premotor cortex (PMd), in the ventral premotor cortex (PMv), and in the right inferior frontal sulcus (IFS) and, subcortically, in the pulvinar, the superior colliculus, and the posterior cerebellum. Clusters and local maxima are listed in Table S1.

**Figure 3.**
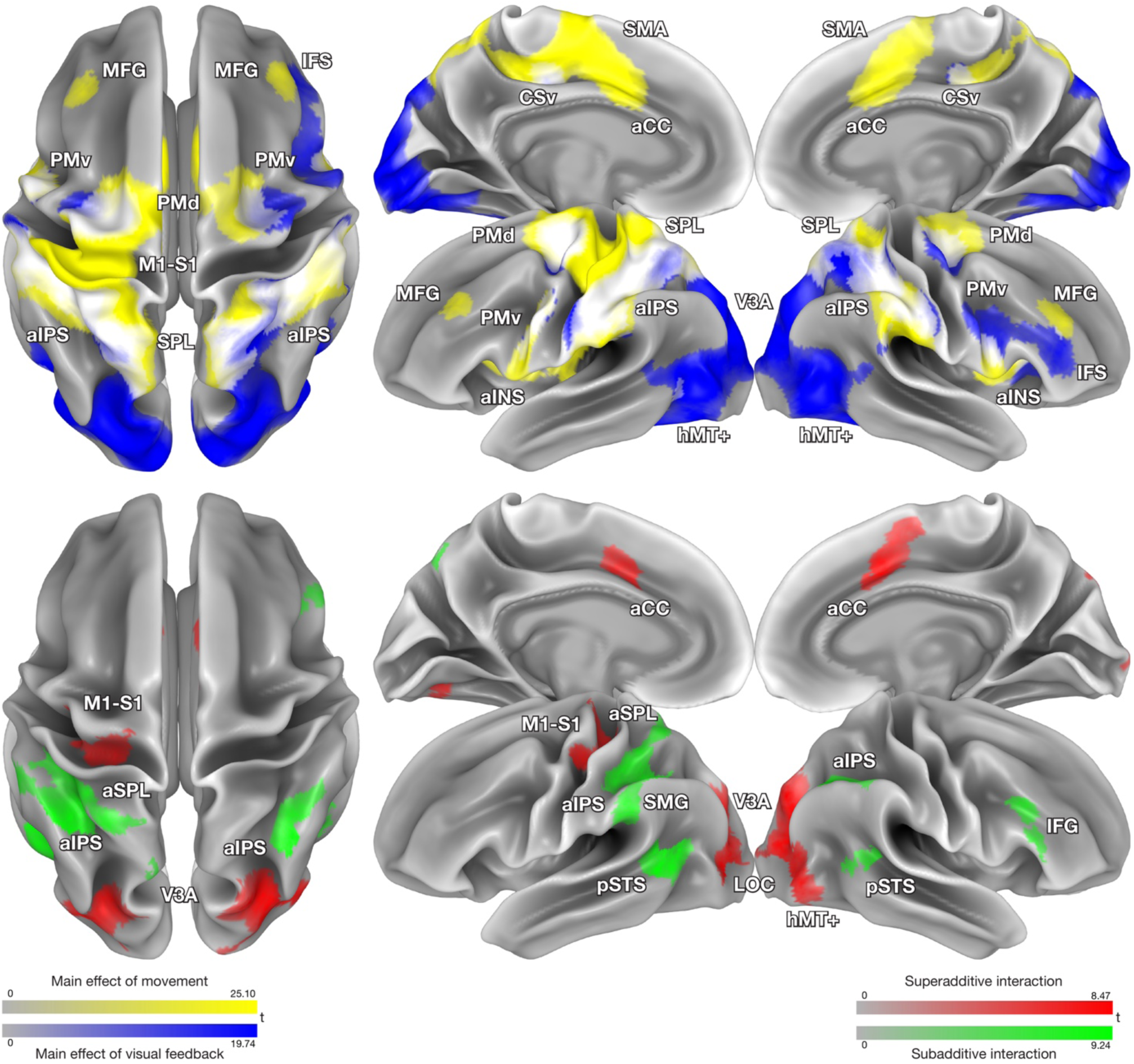
Group-level statistical parametric maps. Top: main effect of movement ([MV + MI] > [SV + SI], yellow) and main effect of hand visual feedback ([MV + SV] > [MI + SI], blue), rendered on the cortical surface in fsLR space. White indicates where the two thresholded maps overlap. Bottom: superadditive ([MV − MI] > [SV − SI], red) and subadditive ([MV − MI] < [SV − SI], green) interactions, masked by the regions significantly activated in either MI or SV relative to SI. Color bars indicate t values, scaled from zero to the peak of each map. All maps were thresholded at *p* < 0.001 at the vertex level and *p* < 0.05 corrected for multiple comparisons at the cluster level through topological FDR. MV, Move Visible; MI, Move Invisible; SV, Stay Visible; SI, Stay Invisible.

The two maps overlapped in a bilateral parieto-frontal network, shown in white in Figure 3, comprising the anterior intraparietal sulcus (aIPS) with the adjacent supramarginal gyrus (SMG), the anterior superior parietal lobule (aSPL), PMd, PMv, and the cingulate sulcus visual area (CSv). Its only occipito-temporal component was a small cluster at the anterior border of the left hMT+. Beyond the cortical surface rendered in Figure 3, the two maps also overlapped in the thalamus and in the posterior cerebellum. Figures S1 and S2 report the response profiles of all regions responding to either factor, expressed as BOLD signal change across the four conditions.

### 3.2 Interactions between movement and hand visual feedback

Subadditive interactions ([MV − MI] < [SV − SI]) — where motor execution combined with visual feedback produced less activity than predicted by motor execution and visual feedback alone — were observed in a bilateral network (Figure 3, green). The largest cluster covered the left aIPS and extended laterally into SMG and dorsally into aSPL. Their activity followed the ordering MV > MI > SV > SI, with both single conditions well above SI (Figure 4), yet the increment obtained by adding visual feedback to movement was smaller than the response to visual feedback alone. In SMG, and in a small cluster in the more medial superior parieto-occipital cortex (SPOC; at coordinates compatible with the parietal reach region: Connolly et al., 2003; Pitzalis et al., 2006), this compression was complete, movement being the dominant drive and visual feedback adding nothing on top of it, if anything slightly reducing it. Notably, in the right aIPS, MV produced no more activity than either single condition. The bilateral pSTS followed a different ordering, SV > MV > MI > SI, indicating that performing the movement that was being seen reduced the response. A cluster in the right inferior frontal sulcus, dorsal and posterior to the one recruited by the main effect of hand visual feedback, showed the same suppression in a more extreme form, responding only during observation without movement and staying at or below baseline in all the other conditions.

**Figure 4.**
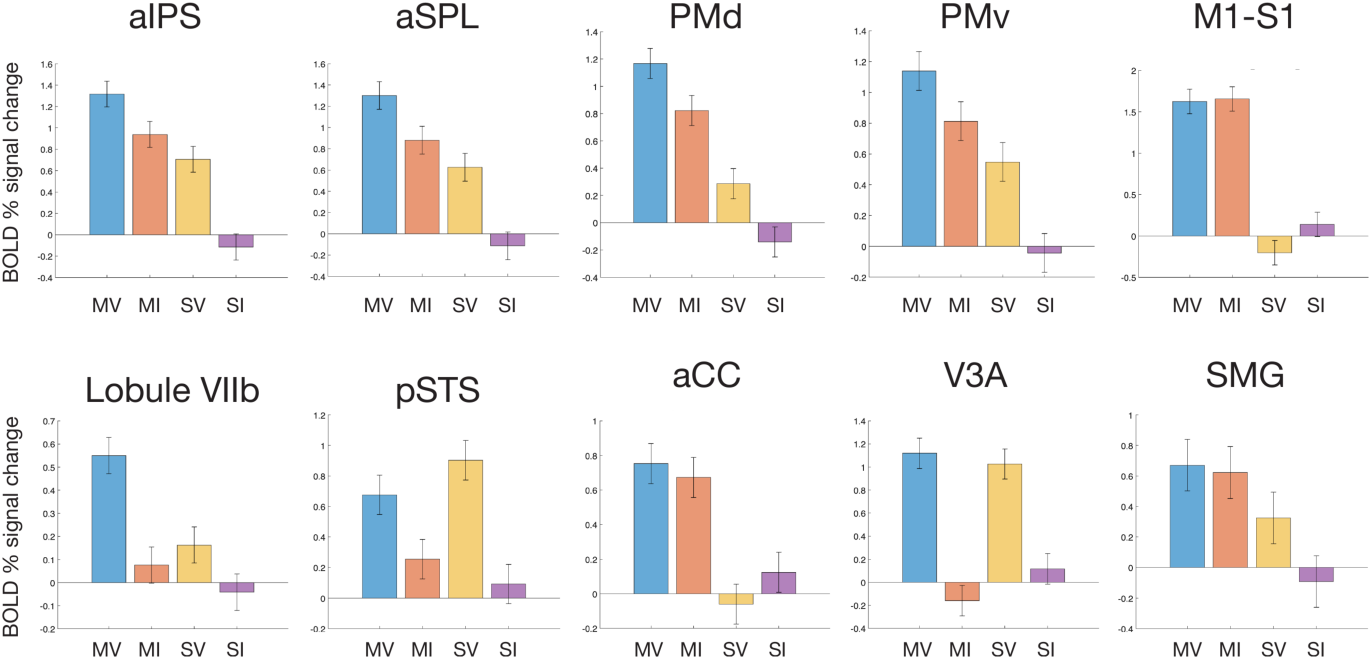
Response profiles across the four conditions. BOLD percent signal change relative to the implicit baseline in the four conditions (MV, Move Visible; MI, Move Invisible; SV, Stay Visible; SI, Stay Invisible), averaged across each cluster, for ten representative regions, all in the left hemisphere. Whiskers represent one standard error of the mean across participants. These values illustrate the shape of the response profiles and were not submitted to further statistical testing, since the same contrasts defined the clusters. Profiles for all the regions responding to either factor and to their interaction are reported in Figures S1—S3.

Superadditive interactions ([MV − MI] > [SV − SI]) — where motor execution combined with visual feedback produced more activity than predicted by motor execution and visual feedback alone (Figure 3, red) — were observed in three sets of regions with distinct response profiles (Figure 4). Large bilateral swaths in the intraoccipital sulcus (IOS/V3A), extending to the lateral occipital complex (LOC) and to hMT+ in the right hemisphere, responded to visual feedback whether participants moved or not, but were suppressed during movement performed in the absence of visual feedback, falling below the level of the SI condition. The left M1-S1 and the bilateral anterior cingulate cortex and anterior insula showed the mirror pattern: they were comparably activated during movement regardless of visual feedback. Still, they were negatively modulated when visual feedback was presented alone. Finally, the posterior cerebellum at the level of lobules VIIb and VIII was the only region in which activity during MV genuinely exceeded the linear combination of activity during both MI and SV, with SI response at baseline. Other cerebellar clusters were found in the vermis and between left lobule VI and Crus I, where movement with visual feedback (MV) produced more activity than movement without visual feedback (MI), and visual feedback per se (SV) was at baseline. See also Figure S3 for the response profiles of all the regions responding to the interaction.

### 3.3 Effective connectivity within the shared network

The connectivity analysis addresses the third hypothesis of the current manuscript, i.e., whether the coupling between the parietal and premotor nodes changes with the availability of visual information about the hand.

To characterize how movement execution and hand visual feedback shape effective connectivity within the reach-to-grasp network, we modeled a network connecting five nodes of the parieto-frontal circuits in the left hemisphere that transform visual and somatosensory information into motor commands, namely aSPL, PMd, aIPS, PMv and M1 (Rizzolatti et al., 1996a,b; Castiello and Begliomini, 2008; Fattori et al., 2010; Battaglia-Mayer and Caminiti, 2019; Bencivenga et al., 2023). All nodes but M1 were chosen because they responded to both movement and hand visual feedback, based on the results of the intersection of the two main effects (shown in white in Figure 3, top row). We added M1 because of its known role in generating movement and defined it using the main effect of movement.

We defined the intrinsic connectivity matrix (matrix A) a priori from known cortico-cortical pathways within and across the parietal and premotor cortex (Rizzolatti et al., 1988; Picard and Strick, 1996; Ferraina et al., 2009; Borra et al., 2017; Caminiti et al., 2017). We modeled a reciprocally connected architecture in which a dorsal pathway (aSPL–PMd) and a ventral pathway (aIPS–PMv) are cross-linked at both the parietal (aSPL–aIPS) and the premotor (PMd–PMv) level, and converge onto M1 through direct connections from PMd and PMv.

Direct driving inputs of conditions onto each region (matrix C) were restricted to a single node per condition, chosen from the univariate results and from the literature before the model was estimated. MV entered at aSPL and SV at aIPS, the parietal nodes most strongly engaged by the main effect of movement and by the main effect of hand visual feedback, respectively. MI entered at PMd, which has been implicated in specifying reach when it cannot be guided by the hand’s visual feedback (Picard and Strick, 1996; Battaglia-Mayer and Caminiti, 2019).

Condition-specific modulations of connectivity (matrix B) were modeled on a subset of the intrinsic connections. MI and MV were allowed to modulate the reciprocal aSPL–PMd and aIPS–PMv connections together with the PMd→M1 and PMv→M1 forward connections, whereas SV, which involves no movement, was allowed to modulate only the reciprocal aSPL–PMd and aIPS–PMv connections, since a change in the gain of the projections onto M1 would have no motor output to act upon.

Figure 5 reports the effective connectivity analysis results: arrows linking the nodes (squares) of the network represent intrinsic connections (matrix A). Modulations of these connections caused by conditions (circles) are depicted as arrows starting from the modeled condition and ending on the connections (matrix B). Driving input to the network is illustrated as an arrow starting from the modeled condition and ending on the modeled node (matrix C). Before either factor acted on the network (intrinsic connectivity, matrix A), the two parietal nodes had positive self-connections, and the three frontal nodes had negative ones. Self-connections in DCM are log-scaled and always inhibitory, so a positive value denotes stronger self-inhibition and a negative value denotes disinhibition (Zeidman et al., 2019a), unlike between-region connections, where a positive value denotes excitation. The parietal end of the network was therefore more strongly self-inhibited, and the frontal end more excitable. Between regions, the coupling was predominantly excitatory. Within the parietal lobe, aSPL and aIPS excited each other, and across lobes aSPL and PMd formed a reciprocally excitatory loop in which the parietal-to-frontal projection was the stronger of the two. At the same time, the model’s two largest parameters ran from aSPL to aIPS and from PMd to M1. PMv was the exception and the only source of inhibition between regions, suppressing aIPS, PMd and, most strongly, M1. It received excitation from PMd and returned inhibition of equivalent size, and M1 weakly excited it. Two of the specified connections did not reach the reporting threshold: the ventral parieto-frontal projection from aIPS to PMv (88.9%) and the feedback from M1 to PMd (60.1%).

**Figure 5.**
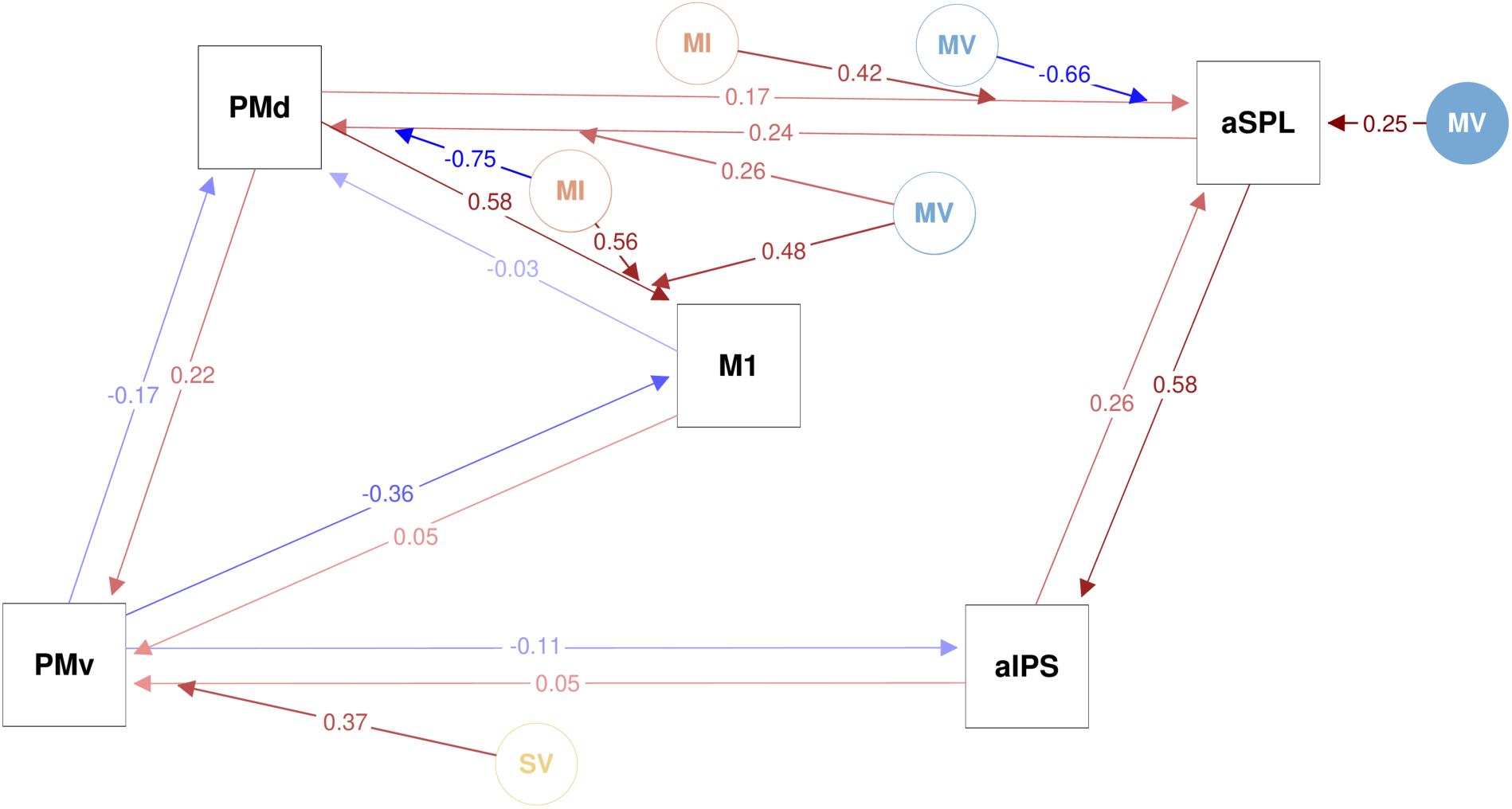
Effective connectivity results. Squares represent the five nodes of the dynamic causal model in the left hemisphere: anterior superior parietal lobule (aSPL), anterior intraparietal sulcus (aIPS), dorsal premotor cortex (PMd), ventral premotor cortex (PMv), and primary motor cortex (M1). Circles represent the experimental conditions that entered the model as driving inputs or as modulatory effects (MV, Move Visible; MI, Move Invisible; SV, Stay Visible). Arrows between squares denote intrinsic effective connectivity (matrix A), arrows from a circle to a square denote driving inputs (matrix C), and arrows from a circle to a connection denote condition-specific modulations of coupling (matrix B). Warm colors indicate excitatory and cool colors indicate inhibitory parameters, with color saturation proportional to effect size. Values are expressed in Hz. Parameters with a posterior probability above 90% are shown, with the addition of intrinsic connections from aIPS to PMv (88.9%) and from M1 to PMd (60.1%). Self-connections are omitted; the full parameter matrices are reported in Figure S4.

For driving inputs (matrix C), only movement with visual feedback, entering at aSPL, drove the network reliably. Movement without visual feedback to PMd was positive but did not reach the reporting threshold, and visual feedback alone to aIPS was retained with a negligible estimate.

Condition-specific changes in coupling (matrix B) fell almost entirely on the reciprocal aSPL–PMd connections, which the two movement conditions modulated in opposite senses. Moving without sight of the hand inhibited the parietal-to-frontal projection from aSPL to PMd and strengthened the frontal-to-parietal feedback from PMd to aSPL, and moving with sight of the hand did the reverse. The coupling present during a condition is the sum of the intrinsic parameter and its modulation, and in each condition the modulation was strong enough to shift one of the two projections from excitatory to inhibitory, not just alter its strength. Excitation moved from the parietal to the frontal cortex when the hand was visible, and from the frontal to the parietal cortex when it was not. The full parameter matrices are reported in Figure S4.

The PMd-to-M1 projection showed a different pattern. Both movement conditions increased it by a similar amount, so the drive to the motor cortex was the same whether the hand was visible or not. What changed with visual feedback was the direction of the excitatory exchange before that output, not the output itself. Seeing the hand move without moving affected another area, exciting the projection from aIPS to PMv, one of the two intrinsic connections that did not reach the reporting threshold, and leaving the dorsal pathway mostly unchanged. In contrast, the two movement conditions had little effect on the ventral connections and the PMv-to-M1 projection. In this model, the dorsal parieto-frontal pathway carries excitation in one direction or the other depending on whether the hand is visible while moving, while seeing the hand alone activates the ventral pathway. We estimated each modulation against zero in a group-mean model, not directly against each other. Therefore, the comparison between conditions is based on two sets of estimates, not a direct contrast.

## 4. Discussion

We orthogonally manipulated movement and the sight of each participant’s moving hand to measure their separate and combined effects. Because participants saw their own movement tracked and replayed, the visual scene was the same in both execution and non-execution conditions. Our three main results match our three hypotheses. First, where the two signals meet in the anterior intraparietal sulcus and anterior superior parietal lobule, their integration is subadditive. Second, in the posterior cerebellum, the integration is superadditive, which fits with internal model updating. Third, the dorsal parieto-frontal pathway carries excitation in opposite directions depending on whether the hand is visible during movement.

### Convergence of movement and hand visual feedback

Both movement and seeing one’s own moving hand activated a broad network, with overlap in the parietal and premotor cortex, which matches the mirroring of observation and execution (Rizzolatti et al., 1996a; Press et al., 2011). Researchers have studied the brain regions involved in movement in both humans and primates, but the effect of seeing one’s own moving hand has only recently gained attention, especially after the discovery that ventral premotor mirror neurons respond to it (Maranesi et al., 2015). The two main effects recruit regions along a gradient: occipito-temporal areas responded to seeing the hand regardless of movement, while the basal ganglia and medial motor regions responded only to movement. In between, in aIPS, aSPL, PMv, and to PMd movement without visual feedback and visual feedback without movement produced similar activity. A convergence zone at the interface of reaching and grasping pathways (Fagg and Arbib, 1998) may help compare predicted and actual hand movements, which is important for online control (Wolpert and Flanagan, 2001), and this matches the mixed signals seen in macaque anterior intraparietal cortex (Lehmann and Scherberger, 2015). Whether these two signals integrate is a separate question, which we address by examining their interaction.

### Subadditivity where prediction meets the signal

The parietal cortex at the level of aIPS and aSPL supported our first hypothesis. Both regions responded to execution without visual feedback and to visual feedback without execution. However, in the left hemisphere, the response to both together was less than the sum of each alone. This is expected for a circuit tuned to the sensory consequences of action. The link between a motor command and seeing the hand move is causal, follows clear rules, and is practiced from infancy, so the circuit encodes the relationship itself rather than each part separately, through repeated co-occurrence and their causal connection. When both signals arrive together, the visual input adds little to what the motor command already predicts. Subadditivity is not a loss of sensitivity but a consequence of efficient coding, since signaling carries a metabolic cost (Attwell and Laughlin, 2001). Thus, an ideal system encodes what it failed to anticipate and not what it already predicts. Optimal feedback control explains this by weighting each source by the inverse of its variance (Diedrichsen et al., 2010a), as do modular models of motor learning (Shadmehr and Mussa-Ivaldi, 1994; Wolpert and Kawato, 1998). The bilateral posterior superior temporal sulcus (pSTS) shows this principle even more strongly: performing a movement that was seen reduced the response below the level seen with visual input alone, meaning a fully predicted visual result is not just explained but reduced (Press et al., 2023). This fits with the connections between the superior temporal region and the parietal and frontal parts of this network (Borra and Luppino, 2017).

### Cerebellar enhancement and model updating

Superadditive responses relate to our second hypothesis, and only one cerebellar cluster met this criterion. At the border between left lobules VIIb and VIII, activity when both the action and its visual result happened together was greater than the sum of each alone, and each alone was close to baseline. A region that responds only to multimodal information is not just comparing single signals, so we do not see it as a comparator. Instead, it appears to be a site of associative plasticity, where pairing an efference copy with a teaching signal creates a sensorimotor mapping (Marr, 1969; Albus, 1971; Ito, 2008), and cerebellar BOLD reflects this input rather than Purkinje cell output (Diedrichsen et al., 2010b). We interpret this as the acquisition of the association, not its use (Kawato et al., 1987; Imamizu et al., 2000). The cluster appears to be where the cerebellum’s second somatomotor area meets its posterior association areas (Buckner et al., 2011; King et al., 2019), where such a mapping would form. Because the cluster is small, we present this as a hypothesis.

### Excitation changes direction in the dorsal parieto-frontal pathway

Most intrinsic connectivity was excitatory and went from the parietal to the frontal cortex along the dorsal pathway, except for PMv, which had inhibitory projections, strongest toward M1. This matches theories where ventral premotor canonical neurons set a motor prototype by balancing inhibition and facilitation over M1 muscle representations (Davare et al., 2007, 2011). The two movement conditions changed the dorsal pathway in opposite ways, with each change strong enough to switch one of its two projections from excitatory to inhibitory. In contrast, just seeing the hand had little effect on this pathway and only influenced a single ventral connection, from aIPS to PMv, which was below threshold.

This reconfiguration was our third hypothesis, which predicted that the coupling would change with the sight of the hand. The parietal-to-frontal projection carries the parietal estimate of the hand’s location, while the frontal-to-parietal projection carries the motor-based prediction of the movement’s sensory effects. When the hand was visible during movement, vision and proprioception together shaped the parietal estimate, and parietal-to-frontal excitation increased. The opposite projection was reduced and became inhibitory. When the hand was not visible, excitation went in the other direction, as if the premotor area had to provide the prediction that vision could not, leaving the parietal area with only proprioception. In this view, the dorsal pathway is where motor predictions meet sensory evidence. Its parietal end, aSPL, is where the response to both signals together was subadditive, while cerebellar superadditivity marks where the predictive model is updated. The aSPL–PMd route supports reaching, and the aIPS–PMv route supports grasping (Fagg and Arbib, 1998; Galletti and Fattori, 2018), so the two parts of prehension (Arbib, 1981; Jeannerod, 1981) seem to be handled differently. This distinction is not absolute (Pisella et al., 2006), and our data show how the two routes are modulated, not what they compute. DCM studies of visually guided grasping have also found that aIPS– PMv coupling changes with the demands of visuomotor transformation (Grol et al., 2007; Begliomini et al., 2015). Together, these analyses identify the circuitry optimized for multimodal information and show how it is integrated across a large network. The factorial interaction pinpoints the circuitry in the anterior parietal regions where movement and the sight of the hand come together, and the effective connectivity in these regions shows how information is carried: bottom-up from parietal to frontal cortex for perception, and top-down in the opposite direction for prediction. Both reflect the predictive brain, and together they show where a cognitive function occurs and how it works.

## Conclusion

By manipulating motor execution and hand visual feedback separately, we could explore how the motor and visual systems work together, not just where each is located. Finding a process and explaining what happens in it are different tasks. A circuit that performs a familiar function may show less activity because it predicts that function well. Activation maps are often interpreted by where the signal is strongest, but the largest signals may reflect prediction errors rather than the circuits that support everyday function. Subadditive interactions may better indicate circuits tuned to a function, suggesting a way to design experiments: combine the signals a circuit is thought to integrate and look for where the response is smaller, not larger. Studying how information is exchanged across the brain, and how this changes with behavior, may reveal important insights.

## Acknowledgments

The authors used Grammarly for grammar and spelling checks. The authors did not use generative artificial intelligence to produce data, analyses, or scientific content.

## Author Contributions

M.T. and G.G. designed the research; M.T., S.F., R.C. and M.P. performed the research; M.T. and G.G. contributed unpublished reagents/analytic tools; M.T., S.F. and G.G. analyzed the data; M.T. and S.F. wrote the first draft of the paper; M.T. and G.G. edited the paper; M.T. and G.G. wrote the paper.

## Supplementary Material

**Figure S1.**
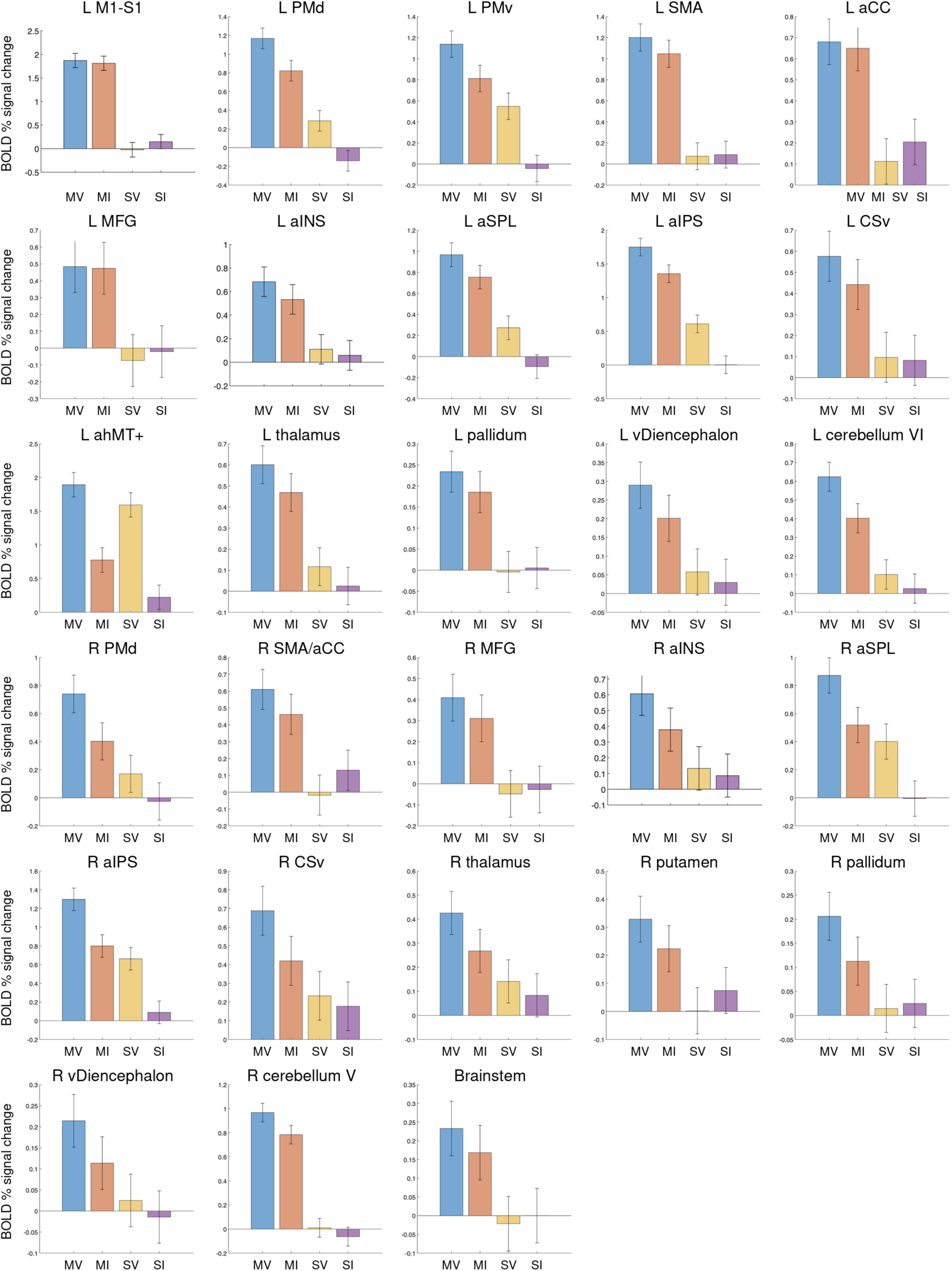
BOLD percent signal change in the regions identified by the main effect of movement ([MV + MI] > [SV + SI]). Each panel shows the mean percent signal change relative to the implicit baseline in the four conditions (MV, Move Visible; MI, Move Invisible; SV, Stay Visible; SI, Stay Invisible), averaged across the cluster surrounding the corresponding local maximum reported in Table S1. Whiskers represent one standard error of the mean across participants. Panels are ordered by hemisphere, left first, and within each hemisphere from primary sensorimotor through premotor, medial frontal, parietal and occipito-temporal cortex to the subcortical structures and the cerebellum. Note the three profiles described in the text: regions responding to execution irrespective of visual feedback, regions showing a graded contribution of both factors, and the anterior portion of the hMT+ complex, in which observation without execution exceeded execution without vision.

**Figure S2.**
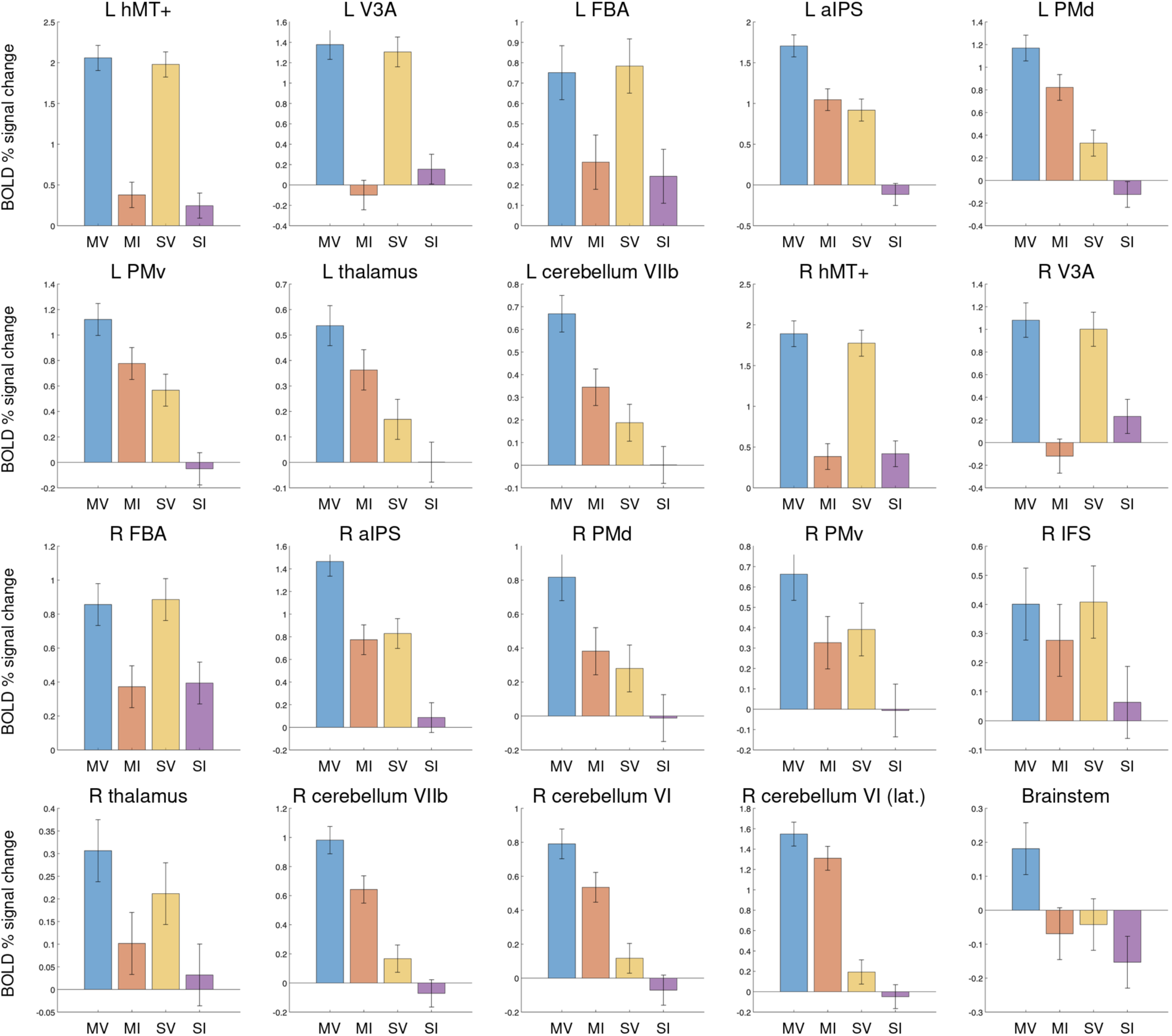
BOLD percent signal change in the regions identified by the main effect of hand visual feedback ([MV + SV] > [MI + SI]). Conventions as in Figure S1, with clusters corresponding to the local maxima reported in Table S1. Occipito-temporal regions show the purely visual profile described in the text, with comparable activity during MV and SV and near-baseline activity whenever visual feedback was absent, whereas the parieto-frontal, thalamic and cerebellar clusters show a contribution of both factors weighted towards movement.

**Figure S3.**
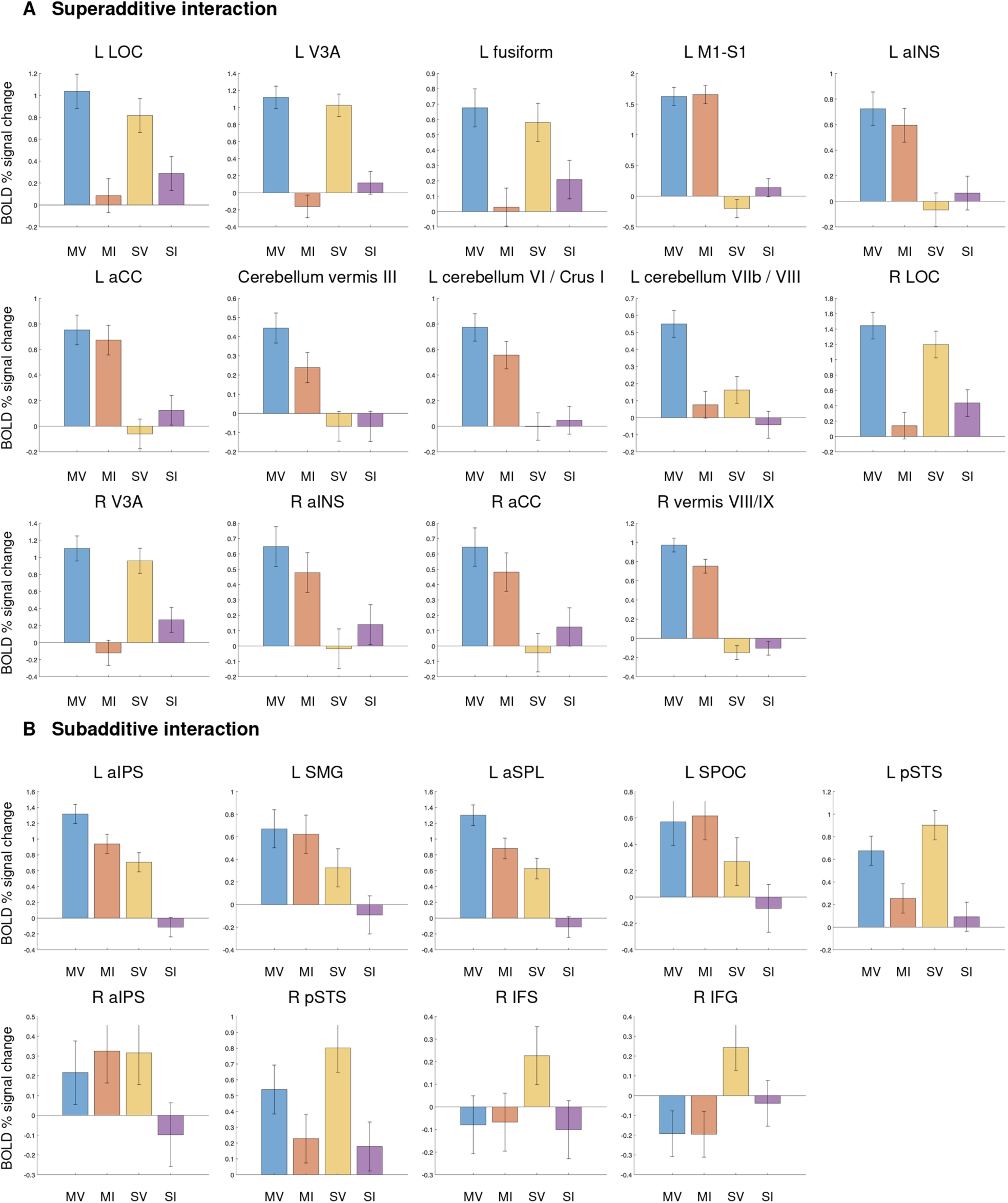
BOLD percent signal change in the regions showing significant interactions between movement and hand visual feedback. Conventions as in Figure S1. A: Regions with a superadditive interaction ([MV -MI] > [SV - SI]), corresponding to the local maxima reported in Table S1. B: Regions with a subadditive interaction ([MV - MI] < [SV - SI]), reported in the same table. The three superadditive profiles and the three subadditive profiles described in the text can be read directly from the panels, from the suppression of the non-preferred modality presented alone to the enhancement of the left posterior cerebellum when visual feedback accompanied self-generated movement, and from the saturation of the left parietal regions to the reduction of the pSTS response when the observed movement was also performed.

**Figure S4.**
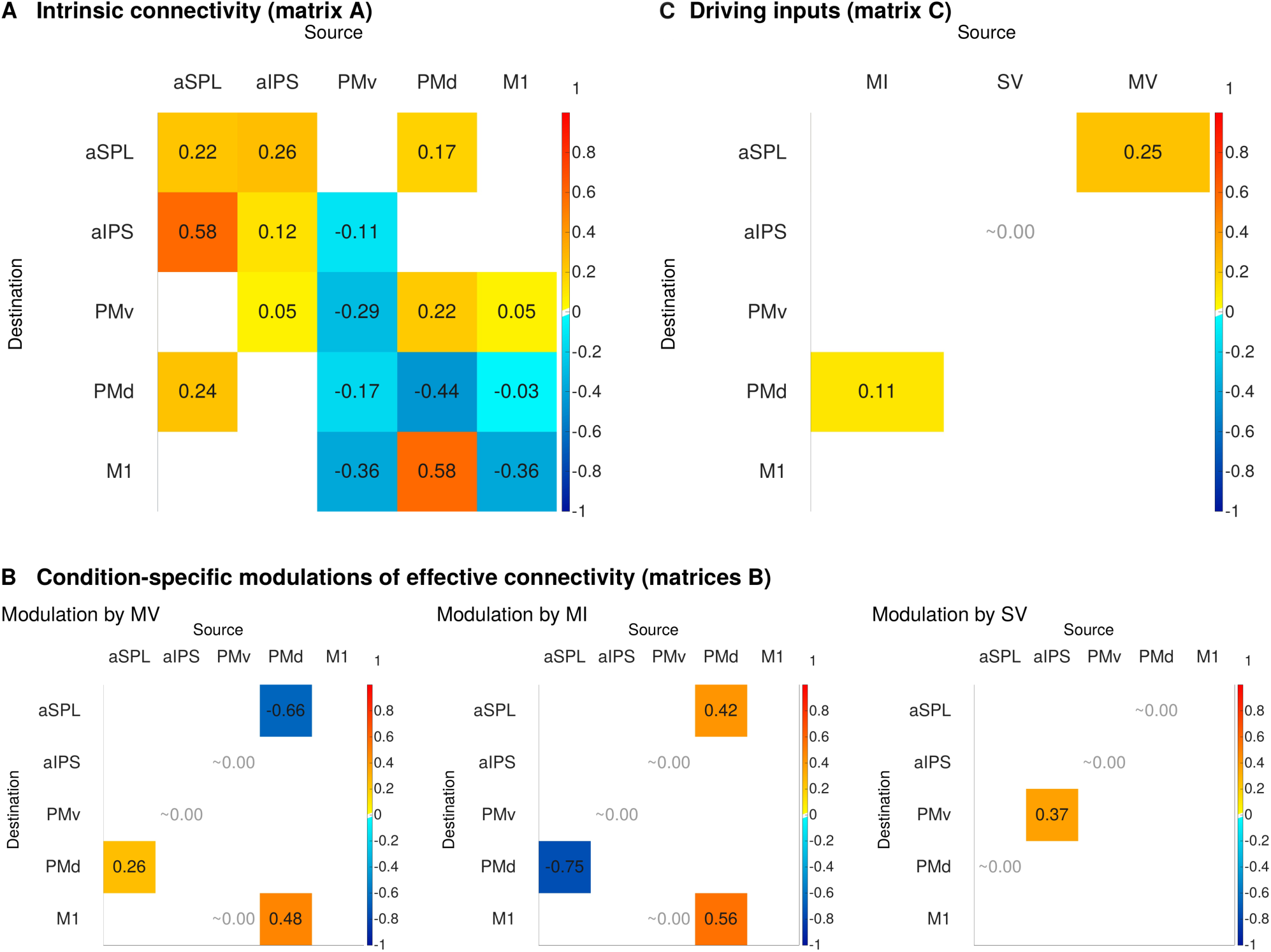
Dynamic causal modeling parameter matrices. A: Intrinsic (average) connectivity, matrix A, showing the coupling between the five nodes in the absence of experimental modulation, that is during Stay Invisible trials and during the periods between the modeled events. B: Condition-specific modulations of effective connectivity, matrix B, one panel per condition. C: Driving inputs, matrix C, showing which condition drives each node directly. Throughout, rows are destinations and columns are sources, values are posterior expectations in Hz for between-region connections and log-scaling parameters for self-connections, for which positive values denote greater self-inhibition and negative values disinhibition. Parameters with a posterior probability above 90% after Bayesian model averaging are displayed, with the addition of intrinsic connections from aIPS to PMv (88.9%), from M1 to PMd (60.1%) and the driving input of movement without visual feedback to PMd (68.4%). Entries marked ∼0.00 were retained with a negligible estimate, and empty entries were pruned. The color scale is common to all panels. MV, Move Visible; MI, Move Invisible; SV, Stay Visible.

**Table S1.**
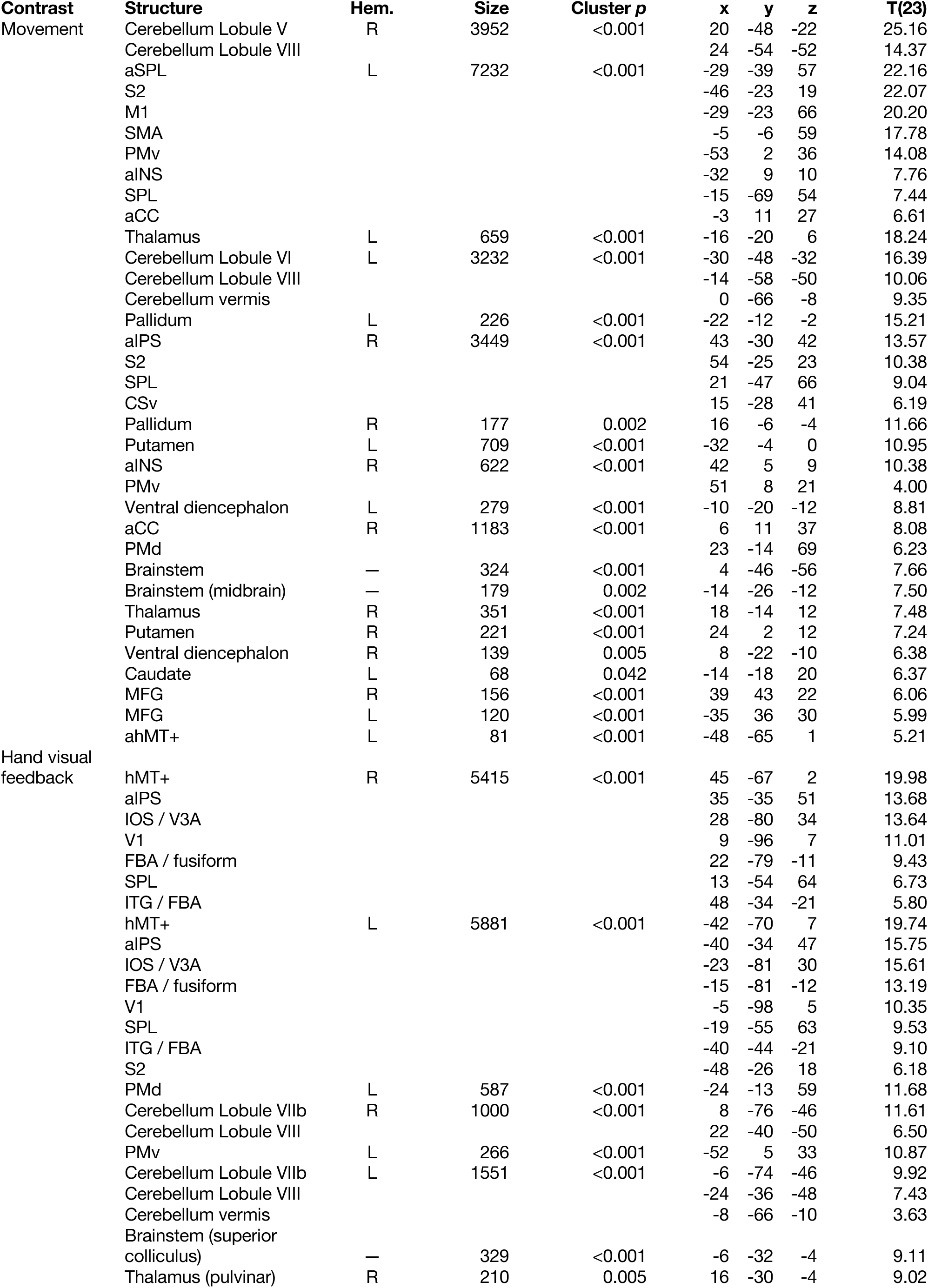
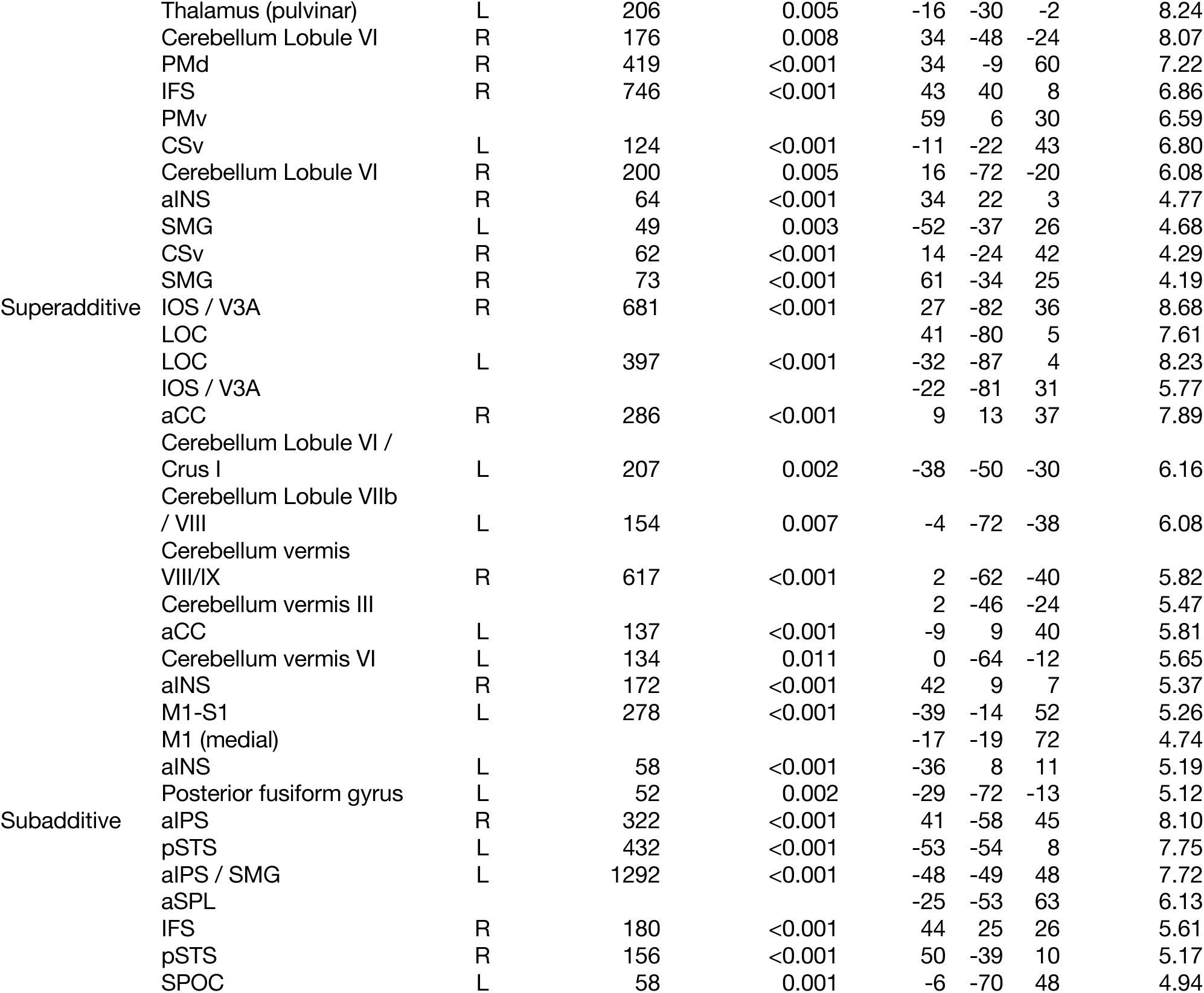
Clusters of significant activation for the main effects of movement and hand visual feedback and for the superadditive and subadditive interactions. Coordinates (x, y, z) are MNI coordinates of the peak vertex or voxel. Hem. = hemisphere (L = left, R = right). Cluster size is expressed in grayordinates, that is surface vertices for cortical clusters and voxels for subcortical and cerebellar clusters. Cluster-level p values are topological FDR-corrected (Chumbley et al., 2010). Rows without a cluster size and cluster p value are local maxima within the cluster listed immediately above, separated by a geodesic distance of at least 40 mm along the cortical surface. T values are reported with 23 degrees of freedom. Clusters are ordered by decreasing peak statistic. Contrasts: movement, [MV + MI] > [SV + SI]; hand visual feedback, [MV + SV] > [MI + SI]; superadditive interaction, [MV − MI] > [SV − SI]; subadditive interaction, [MV − MI] < [SV − SI]. Abbreviations: aCC, anterior cingulate cortex; aINS, anterior insula; aIPS, anterior intraparietal sulcus; aSPL, anterior superior parietal lobule; CSv, cingulate sulcus visual area; FBA, fusiform body area; ahMT+, anterior portion of the human middle temporal complex; hMT+, human middle temporal complex; IFS, inferior frontal sulcus; IOS, intraoccipital sulcus; ITG, inferior temporal gyrus; LOC, lateral occipital complex; M1, primary motor cortex; MFG, middle frontal gyrus; PMd, dorsal premotor cortex; PMv, ventral premotor cortex; pSTS, posterior superior temporal sulcus; S1, primary somatosensory cortex; S2, secondary somatosensory cortex; SMA, supplementary motor area; SMG, supramarginal gyrus; SPL, superior parietal lobule; SPOC, superior parieto-occipital cortex; V1, primary visual cortex; V3A, visual area V3A.

